# Mechanisms of mitochondrial reactive oxygen species action in bone mesenchymal cells

**DOI:** 10.1101/2025.03.24.643319

**Authors:** Md Mohsin Ali, Intawat Nookaew, Ana Resende-Coelho, Adriana Marques-Carvalho, Aaron Warren, Qiang Fu, Ha-Neui Kim, Charles A O’Brien, Maria Almeida

## Abstract

Mitochondrial reactive oxygen species (mtROS), insufficient NAD^+^, and cellular senescence all contribute to the decrease in bone formation with aging. ROS can cause senescence and decrease NAD^+^, but it remains unknown whether these mechanisms mediate the effects of ROS *in vivo*. Here, we generated mice with deletion of the mitochondrial antioxidant enzyme Sod2 in Osx1-Cre (Sp7-tTA, tetO-EGFP/cre) targeted cells designated *Sod2*^Δ*Osx1*^ mice. We showed that Sod2 deletion caused low bone mass. Osteoblastic cells from these mice had impaired mitochondrial respiration and attenuated NAD^+^ levels. Administration of an NAD^+^ precursor improved mitochondrial function *in vitro* but failed to rescue the low bone mass of *Sod2*^Δ*Osx1*^ mice. Single-cell RNA-sequencing of bone mesenchymal cells indicated that ROS had no significant effects on markers of senescence but disrupted parathyroid hormone signaling, iron metabolism, and proteostasis. Our data support the rationale that treatment combinations aimed at decreasing mtROS and senescent cells and increasing NAD^+^ should confer additive effects in delaying age-associated osteoporosis.

## Introduction

Aging is a major cause of osteoporosis and increased fracture risk (1–3). Age-related bone loss is primarily due to a decline in bone formation caused by a reduced number of osteoblasts — the cells responsible for the secretion of bone matrix (4). Osteoblasts are short-lived cells that originate from mesenchymal precursors. We have previously shown that ROS increases in bone with aging and that suppression of mitochondrial ROS in all cells of the mesenchymal lineage, including stromal cells, osteoblasts precursors, osteoblasts, and osteocytes, attenuates age-associated bone loss (3, 5). In line with the damaging role of ROS in bone during aging, deletion of the antioxidant enzyme *Sod2* in cells of the osteoblast lineage in mice is sufficient to reduce bone formation and decrease bone mass in young adult mice (6, 7).

The majority of cellular ROS are generated in the mitochondria during oxidative phosphorylation due to the leakage of electrons passing through the electron transport chain (8, 9). Electrons react with molecular oxygen to generate superoxide radicals (O2•–) which are rapidly converted to hydrogen peroxide (H_2_O_2_) by Cu/Zn-superoxide dismutase (Cu/ZnSOD or SOD1) and manganese-dependent superoxide dismutase (MnSOD or SOD2) (9). One of the known pathological effects of O_2_^•–^ is the disruption of the Fe-S clusters in aconitase, a critical tricarboxylic acid (TCA) cycle enzyme (10, 11). Excessive ROS can also decrease the NAD^+^/NADH redox balance, which is essential to cellular metabolism. Indeed, NADH is the major electron donor to complex I. NAD^+^ is reduced back to NADH during multiple metabolic reactions but is also utilized by many NAD-dependent enzymes, including the poly (ADP-ribose) polymerases (PARPs) and sirtuin family of proteins (12). We have shown previously that NAD^+^ decreases with aging in osteoblastic cells and administration of NAD^+^ precursors to mice attenuates the decrease in bone mass with aging (13).

Excessive ROS can also cause cellular senescence, and an increase in osteoblastic cell senescence contributes to the deficient bone formation and loss of bone mass with aging (14–16). Senescent cells exhibit cell cycle arrest due to the upregulation of p16 (*Cdkn2a*) or p21 (*Cdkn1a*) (17). Other changes in senescent cells include high senescence-associated β-galactosidase (SA-β-gal) activity, dysfunctional mitochondria, and secretion of an array of pro-inflammatory cytokines, chemokines, and proteases, known collectively as the senescence-associated secretory phenotype (SASP) (18, 19). SASP factors can either reinforce growth arrest in an autocrine manner or cause senescence in surrounding cells in a paracrine manner.

While an increase in ROS is a well-established cause of skeletal aging, the mechanisms that mediate the deleterious effects of ROS on bone formation remain unclear. It is also unknown whether excessive ROS in osteoblastic cells is causally related to other mechanisms of aging, such as senescence and decreased NAD^+^. Understanding possible relationships between different mechanisms might contribute to the development of optimal treatment combinations to combat the loss of bone mass with aging.

Herein, we examined whether excessive levels of mtROS, such as that caused by *Sod2* deletion, are sufficient to cause other age-associated dysfunctions, such as cellular senescence and NAD^+^ deficiency, and whether these effects could mediate the loss of bone mass due to mtROS. We also used the unbiased approach of single-cell RNA-sequencing (scRNA-seq) to identify potential mechanisms via which excessive mtROS affects mesenchymal bone cells.

## Results

### Deletion of *Sod2* in osteoblast lineage cells decreases bone mass

To investigate the effect of mtROS in cells of the osteoblast lineage, we generated mice in which cells targeted by an *Osx1-Cre* transgene were deficient in superoxide dismutase 2 (*Sod2*), referred to as *Sod2*^Δ*Osx1*^ mice. This was accomplished by crossing *Osx1-Cre* (20) with *Sod2 floxed (^f/f^)* mice (21) (**Fig. S1A**). *Osx1-Cre* littermates were used as controls. Bone marrow-derived stromal cell (BMSC) cultures of *Sod2*^Δ*Osx1*^ mice exhibited a decrease in *Sod2* mRNA levels, confirming effective deletion of the gene in the osteoblast lineage (**Fig. 1A**). 13-week-old female *Sod2*^Δ*Osx1*^ mice had lower bone mineral density (BMD) both at the lumbar spine and femur, as determined by dual-energy X-ray absorptiometry (**Fig. 1B**). Likewise, male *Sod2*^Δ*Osx1*^ mice had reduced BMD at both sites (**Fig. 1B**). Examination of bone microarchitecture by micro-computed tomography (*µ*CT) revealed that cortical thickness, measured at the femur midshaft, was lower in 26-week-old male *Sod2*^Δ*Osx1*^ mice (**Fig. 1C**). Trabecular bone volume at the fifth lumbar vertebra (L5) was also lower in *Sod2*^Δ*Osx1*^ mice (**Fig. 1D**). The lower bone volume was due to a decrease in trabecular number and was associated with an increase in trabecular separation. Trabecular thickness was not affected by *Sod2* deletion (**Fig. S1B**). Dynamic histomorphometry showed that mineralizing surface (MS/BS), mineral apposition rate (MAR), and bone formation rate (BFR/BS) were not different between control and *Sod2*^Δ*Osx1*^ mice as determined at the lumbar vertebrae (**Fig. S2A**). Likewise, serum bone turnover markers (PINP and CTX) were unchanged (**Fig. S2B**). Most likely, the cellular changes that led to the low bone mass phenotype occurred at an earlier age. Overall, these results demonstrate that deletion of *Sod2* in the osteoblast lineage is sufficient to decrease bone mass in young adult mice and confirm previous studies in which the gene was deleted with *Dmp1-Cre* and *Runx2-Cre* (6, 7).

**Figure 1.**
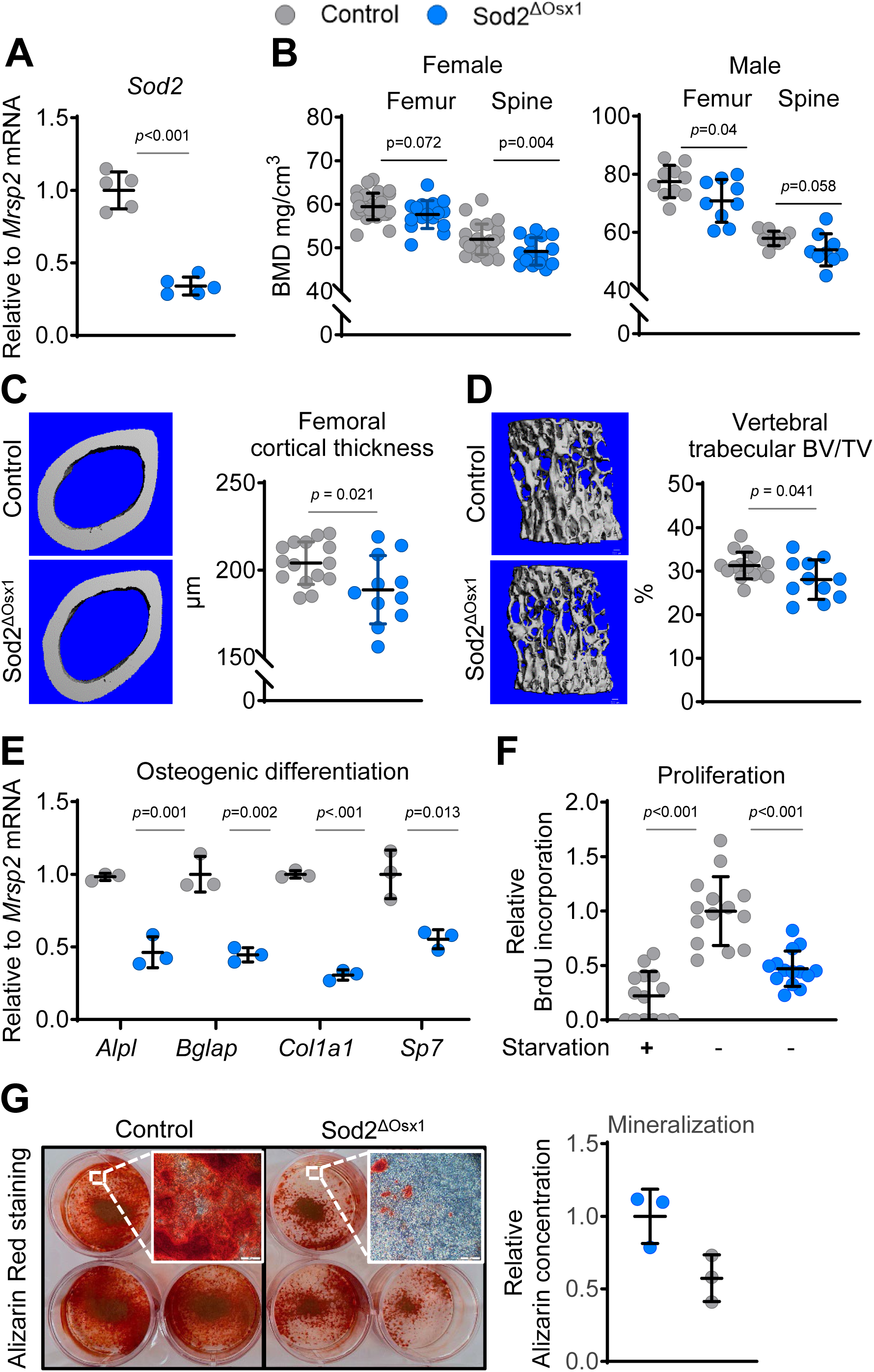
Deletion of *Sod2* in osteoblast lineage cells causes low bone mass. (A) Quantitative RT-PCR analysis of *Sod2* mRNA levels in BMSCs cultured for 4 days (biological replicates *n* = 5/group). (B) Bone-mineral density (BMD) in 13-week-old females (left; femur and lumbar spine; Control n = 23-26, *Sod2*^Δ*Osx1*^ n = 15-16 mice) and 16-week-old males (left; femur and lumbar spine; Control n =10, *Sod2*^Δ*Osx1*^ n = 9). (C and D) Representative μCT images (left) and quantification (right) of (C) femoral cortical thickness and (D) L5 vertebral trabecular BV/TV from 26-week-old male mice (n = 9-10 mice/group). Also see Figure S1 (E) mRNA expression levels of osteoblast marker genes in BMSCs cultured for 7 days in osteogenic medium (technical replicates, n = 3/group). (F) BrdU-incorporation assay in BMSCs cultured for 48 h with 10 µM BrdU; serum-starved BMSCs for 24 h served as a negative control (technical replicates: Neg Ctrl n = 8, Control n = 14, *Sod2*^Δ*Osx1*^ n = 14). (G) Alizarin Red S staining of BMSCs cultured for 21 days in osteogenic medium (representative images; inset scale bar = 500 µm) with quantification (technical replicates, n = 3 per group). Data are meansD±DSD. P values by two-tailed unpaired Student’s t-test. Each technical replicate consisted of BMSCs pooled from 3–6 mice/group.

The mRNA levels of osteoblastic marker genes such as *Alpl*, *Bglap*, *Col1a1*, and *Sp7* were decreased in BMSCs derived from *Sod2*^Δ*Osx1*^ mice following 7 days of osteogenic differentiation (**Fig. 1E**). BrdU incorporation was also reduced in these cells (**Fig. 1F**). In line with these results, Alizarin Red S staining of the calcified matrix was substantially lower in cells with *Sod2* deletion (**Fig. 1G**). These findings suggest that a decrease in osteoblast number contributes to the low bone mass caused by *Sod2* deletion, as shown by others using similar mouse models (6, 7).

### Deletion of *Sod2* causes mitochondrial dysfunction mimicking the effects of aging in BMSCs

We next examined the effects of *Sod2* deletion on mitochondrial biology. As anticipated, mtROS levels were elevated in BMSCS from *Sod2*^Δ*Osx1*^ mice compared to the control group, indicated by MitoSOX measurements (**Fig. 2A**). Additionally, mitochondrial membrane potential (ΔΨm), crucial for mitochondrial function, was decreased with *Sod2* deletion (**Fig. 2B**). To evaluate mitochondrial respiration, we measured the oxygen consumption rate (OCR) in BMSCs. Both basal and maximal respiration were lower in cells from *Sod2*^Δ*Osx1*^ mice (**Fig. 2C**). Similar effects on mitochondria were seen in BMSCs cultured from old versus young wild-type mice. Specifically, cell cultures from 104-week-old C57BL/6 mice had higher mtROS (**Fig. 2D**), reduced ΔΨm (**Fig. 2E**), and lower basal and maximal respiration (**Fig. 2F**) when compared to cells from 26-week-old mice of the same strain. These results support the idea that deletion of *Sod2* causes a dysfunction in mitochondria similar to that seen in old mice.

**Figure 2.**
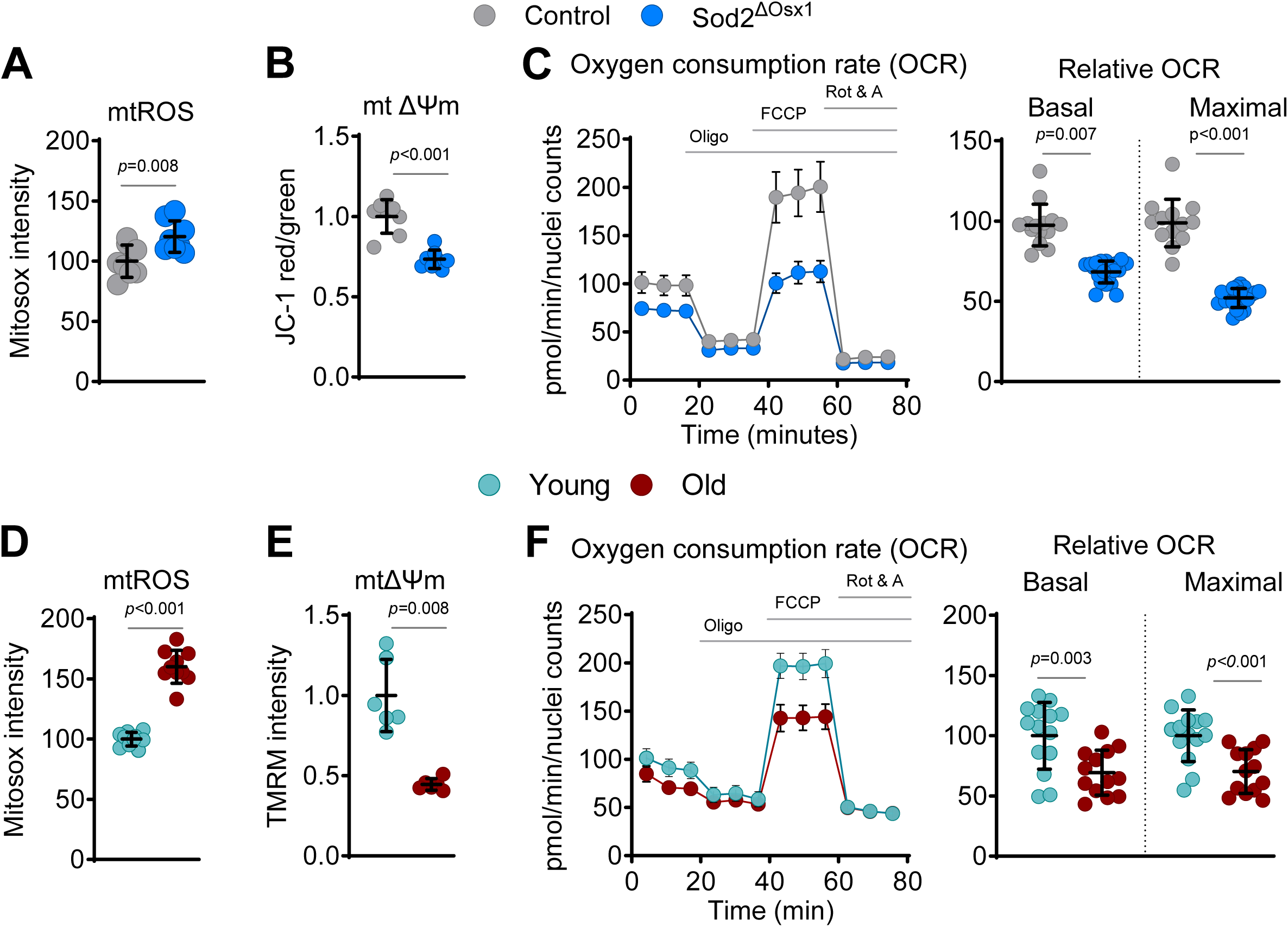
Deletion of *Sod2* causes mitochondrial dysfunction mimicking the effects of aging in BMSCs. (A and D) Mitochondrial ROS (mtROS) levels in cultured BMSCs detected using the fluorescent probe 2.5 µM MitoSOX. (A) Control vs. *Sod2*^Δ*Osx1*^ (technical replicates, n = 8/group). (D) Young vs. old (technical replicates, n = 10). (B and E) Mitochondrial membrane potential (ΔΨm) measured in cultured BMSCs (B) Control vs. *Sod2*^Δ*Osx1*^, measured with 2 µM JC-1; a lower red/green ratio indicates depolarization (technical replicates, n = 8/group). (E) Young vs. old, measured with 100 nM TMRM; lower fluorescence denotes reduced ΔΨm (technical replicates, n = 6). (C and F) Oxygen-consumption rate (OCR) in live BMSCs, with basal and maximal mitochondrial respiration quantified. (C) Control vs. *Sod2*^Δ*Osx1*^ (technical replicates, basal n = 14/group, maximal n = 20/group). (F) Young vs. old (technical replicates n = 13-14). Data are meansD±DSD. P values by a two-tailed unpaired Student’s t-test. Each technical replicate consisted of BMSCs pooled from 3–6 mice/group.

### NR treatment increases NAD^+^ and mitochondrial respiration in osteoblastic cells in culture

Elevated mtROS can decrease the levels of NAD^+^ by impairing respiratory complex I and the oxidation of NADH to NAD^+^ and by stimulating DNA repair enzymes that consume NAD^+^ (22, 23). Cellular NAD^+^ levels can be replenished by treatment with the NAD^+^ precursor nicotinamide riboside (NR). We found that cultured *Sod2*-deficient BMSCs had lower NAD^+^ than control cells (**Fig. 3A**). Addition of NR to the cultures modestly increased NAD^+^ in control cells and restored the basal levels of NAD^+^ in cells from *Sod2*^Δ*Osx1*^ mice. We also assessed mitochondrial function by measuring OCR following 16 hours of NR treatment (**Fig. 3B**). NR increased basal respiration in cells from *Sod2*^Δ*Osx1*^ mice. An increase in maximal respiration was seen in cells from both genotypes but was more robust in cells from *Sod2*^Δ*Osx1*^ mice (**Fig. 3C**). These results demonstrated that NR supplementation attenuates the damaging effects of *Sod2* deletion in cells of the osteoblastic lineage and suggested that it might alleviate the negative effect on the skeleton.

**Figure 3.**
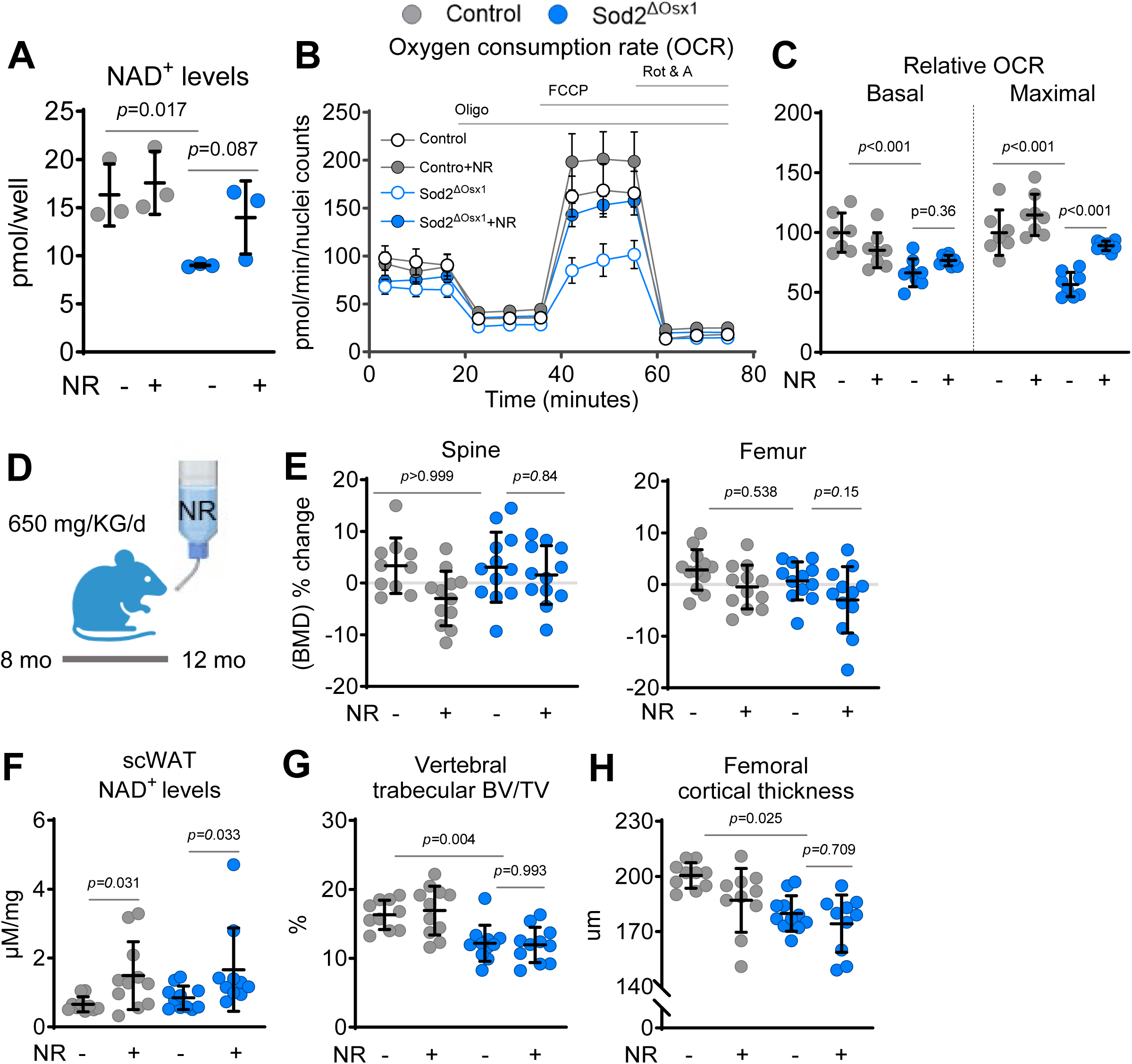
NR stimulates mitochondrial respiration in osteoblastic cell cultures but does not affect bone mass of *Sod2*^Δ*Osx1*^ mice. (A) Cellular NAD^+^ levels in cultured BMSCs from control and *Sod2*^Δ*Osx1*^ mice (technical replicates, n = 4/group). (B, C) OCR in BMSCs after 16 h treatment with water (control) or 5mM NR. Oligo (oligomycin, an ATP synthase inhibitor), FCCP (carbonyl cyanide-4 (trifluoromethoxy) phenylhydrazone, an uncoupler of oxidative phosphorylation), and Rot & A (rotenone and antimycin A, inhibitors of complex I and III, respectively) were sequentially added to the cultures as indicated. (C) Quantification of basal and maximal respiration from the assays in (B) (technical replicates n = 7–8/group). BMSCs pooled from 3–6 mice/group (D) Schematic of *in vivo* NR supplementation in control and *Sod2*^Δ*Osx1*^ mice. (E) Percentage change of lumbar spine and femoral BMD (n = 10-12 mice/group). (F) Quantification of NAD^+^ in subcutaneous white adipose tissue (scWAT) (n = 10-11 mice/group). (G and H) μCT analysis in (G) L5 lumbar vertebrae and (H) femur (n = 10-11 mice/group). See also Figure S2. Data are meanD±DSD. P-values by two-way ANOVA with Tukey’s multiple comparisons test.

### NR supplementation does not rescue the low bone mass in *Sod2*^Δ^*^Osx1^* mice

We next administered NR in the drinking water to 35-week-old female control and *Sod2*^Δ*Osx1*^ mice for 17 weeks to evaluate its effects on bone mass (**Fig. 3D**). We found that NAD^+^ levels were increased in subcutaneous white adipose tissue (scWAT) of NR-treated mice of both genotypes (**Fig. 3F**). Nevertheless, this intervention did not alter BMD in the lumbar spine or femur, as both regions exhibited no significant changes compared to untreated controls (**Fig. 3E**). µCT analysis revealed that female *Sod2*^Δ*Osx1*^ mice, similar to male, had decreased trabecular bone volume (BV/TV) in L5 vertebra (**Fig. 3G**) due to reduced trabecular number (**Fig. S3A**) and lower femoral cortical thickness (**Fig. 3H**) due to smaller cortical area, and the ratio of cortical area to total cross-sectional area within the periosteal envelope (**Fig. S3B**). However, NR supplementation did not affect any of these parameters in the lumbar vertebrae or femur from either *Sod2*^Δ*Osx1*^ or control mice. Thus, while NR reduced signs of mitochondrial damage in osteoblast lineage cells, this was not sufficient to counteract the negative effects of *Sod2* deletion.

### scRNA-seq reveals that *Sod2* deletion greatly changes the transcriptome of osteoblasts and adipo-CAR

To examine the transcriptional changes induced by elevated ROS in osteoblast lineage cells, we harvested endosteal cells from 19-week-old male *Sod2*^Δ*Osx1*^ and littermate control mice. The endosteal cells were isolated from tibias and femurs after removing the periosteum, epiphyses, and bone marrow, followed by sequential incubations with collagenase and EDTA, as previously described (24) (**Fig. 4A**). A total of 15,224 cells were subjected to scRNA-seq using the 10X Chromium platform. Clustering analysis using the Louvain algorithm revealed distinct clusters representing major mesenchymal cell types, with a small proportion of hematopoietic and endothelial cells identified based on established markers (**Fig. 4B, S3A**). The major mesenchymal cell types present at the endosteum were osteoblasts, pre-osteoblasts, osteo-Cxcl12 abundant reticular (CAR), and adipo-CAR, as previously described (24). Osteoblasts were defined by high expression of osteocalcin (*Bglap*), pre-osteoblasts by osteopontin (*Spp1*), osteo-CAR by LIM and calponin homology domains 1 (*Limch1)*, adipo-CAR were identified by expression of C-X-C motif chemokine ligand 12 (*Cxcl12*), and osteocytes were identified by phosphate regulating endopeptidase X-linked (*Phex*) (**Fig. 4C**). Although osteocytes are abundant in bone, they were not obtained in large numbers, likely due to their inefficient release from calcified bone matrix. It has been previously proposed that osteocytes mediate the deleterious effects of *Sod*2 deletion on bone mass (7). However, due to the low numbers obtained, we could not perform a reliable analysis of the transcriptome of osteocytes lacking *Sod2*.

**Figure 4.**
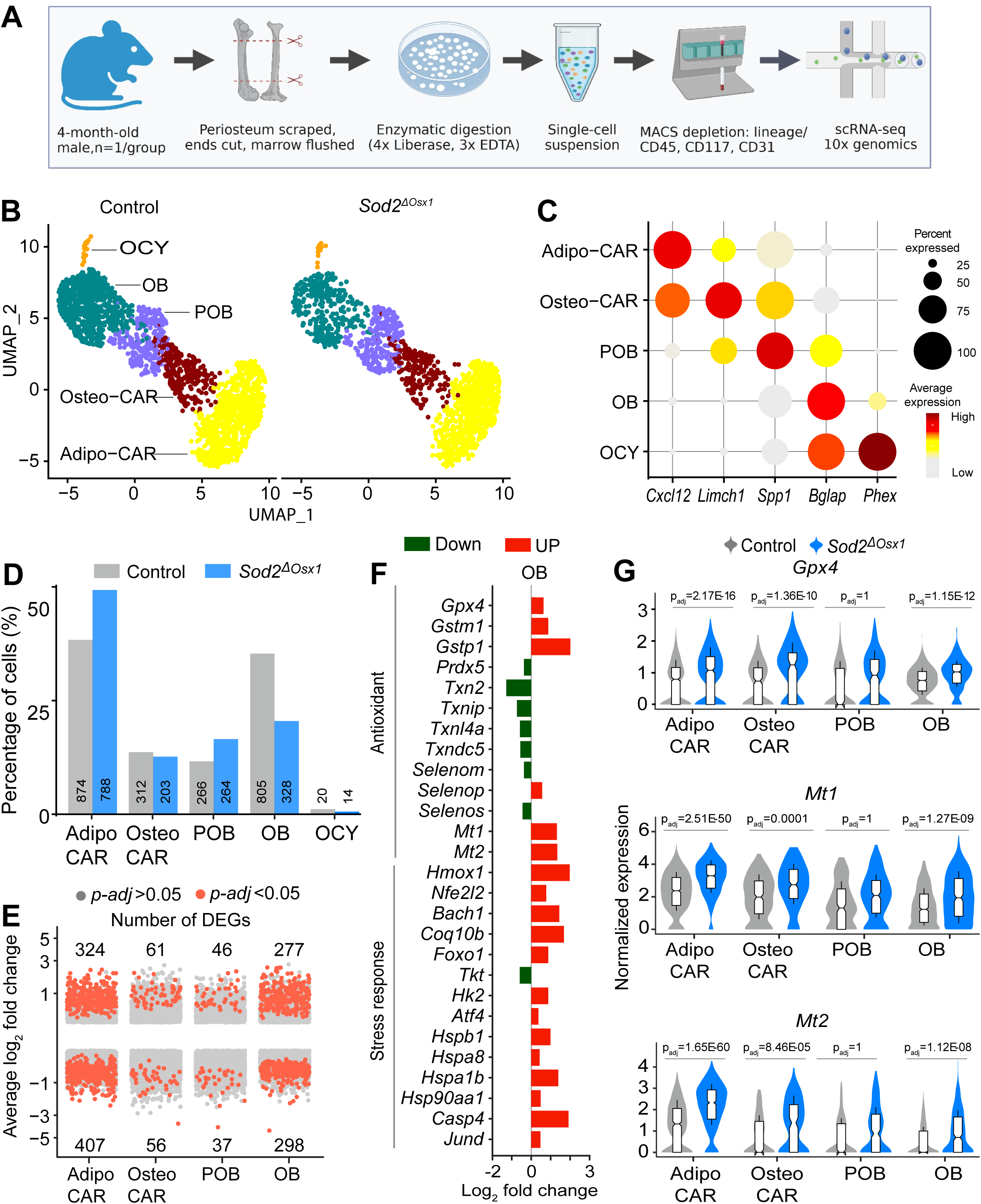
scRNA-seq reveals that *Sod2* deletion greatly impacts the transcriptome of osteoblasts and adipo-CAR. (A) Schematic of endosteal mesenchymal cell isolation from femurs and tibias of 4-month-old male *Sod2*^Δ*Osx1*^ and control mice (n=1/group) enriched in mesenchymal lineage cells and subjected to scRNA-seq using the 10X Chromium platform. (B) Uniform Manifold Approximation and Projection (UMAP) of endosteal mesenchymal cell clusters: Adipo-CAR, Osteo-CAR, preosteoblast (POB), osteoblast (OB), and osteocytes (OCY) identified by scRNA-seq. Each dot represents a single cell, color-coded by clusters. See Figure S3. (C) Dot plot showing the expression of marker genes (*Cxcl12*, *Limch1*, *Spp1*, *Bglap*, *Phex*) across cell types. Dot size indicates the percentage of cells expressing the gene; color intensity represents the average expression level. (D) Comparison of specific cell cluster proportions between *Sod2*^Δ*Osx1*^ and control mice. Numbers below the bars indicate the percentage of each cluster for each group. (E) Distribution and directionality of differentially expressed genes (DEGs). Significant DEGs (p-adj < 0.05, red dots), with numbers of up/downregulated DEGs, are indicated above/below bars for each cell cluster. (F) Log_2_ fold change (Log_2_FC) of antioxidant and stress response genes in OB clusters from *Sod2*^Δ*Osx1*^ compared to control. (G) Violin plots of normalized expression levels of Gpx*4*, *Mt1* and *Mt2* in Adipo-CAR, Osteo-CAR, POB, and OB clusters from *Sod2*^Δ*Osx1*^ compared to control. Data are box-and-whisker plots within the violins, (center line: median; box limits: 25th/75th percentiles; whiskers ±1.5× interquartile range).

We first examined whether elevated ROS levels are associated with changes in the relative abundance of any of the clusters. Osteoblasts exhibited the most significant reduction in abundance with elevated ROS, decreasing from 35.3% to 20.5% (**Fig. 4D**). This reduction is consistent with previous histomorphometric findings indicating that osteoblast number decreases upon *Sod2* deletion (25). To identify changes in gene expression associated with elevated ROS in *Sod2*^Δ*Osx1*^ mice compared to controls, we performed differential expression analysis for each cell cluster using Model-based Analysis of Single-cell Transcriptomics (MAST) with a false discovery rate (FDR)D<D0.05 and a random effect for sample origin (26). In osteoblasts, we identified 575 differentially expressed genes (DEGs) (adjusted pD<D0.05), among which 277 were upregulated and 298 downregulated (**Fig. 4E**). Similarly, in adipo-CAR cells, we found 731 DEGs (adjusted pD<D0.05), with 324 upregulated and 406 downregulated. Both pre-osteoblast and osteo-CAR exhibited fewer than 117 DEGs. As anticipated, cells from *Sod2*^Δ*Osx1*^ mice had altered oxidant detoxification genes. For example, glutathione peroxidase 4 (*Gpx4*), and glutathione S-transferases *Gstm1* and *Gstp1*, which detoxify lipid hydroperoxides, were all upregulated, as were metallothionein 1 and 2 (*Mt1 and Mt2*) (**Fig. 4F, 4G**). Multiple transcription factors implicated in the response to oxidative stress, including BTB and CNC homology 1 *(Bach1*), Nuclear factor erythroid 2-related factor 2 (Nrf2/*Nfe2l2*), forkhead box O1 (*Foxo1*), and activating transcription factor 4 (*Atf4*) were also increased. In contrast, thioredoxins and ER-resident selenoproteins were downregulated (**Fig. 4F**).

### *Sod2* deletion alters biological processes related to iron homeostasis and translation but not cellular senescence, in all major mesenchymal cell clusters

To identify biological processes impacted by *Sod2* deletion, we performed Gene Ontology (GO) enrichment analysis(27) (**Fig. 5A**). Superoxide can damage proteins containing FeS clusters, such as aconitase, causing the release of ferrous iron and altered citrate metabolism (28, 29). Genes associated with iron sequestration and detoxification were upregulated, including the ferritin subunit *Ftl1* (**Fig. 5B, 5C**) and heme oxygenase 1 (*Hmox1*), likely to mitigate Fenton-driven oxidative damage (**Fig. 5B**). Genes related to rRNA processing, cytoplasmic translation, and mTOR signaling were also upregulated. These included genes encoding translation initiation factors such as *Eif1, Errfi1,* and *eIF4E* and numerous ribosomal proteins (**Fig. 5B**). Genes related to several other cellular stress markers were upregulated, including angiogenesis, p38MAPK and p53 signaling.

**Figure 5.**
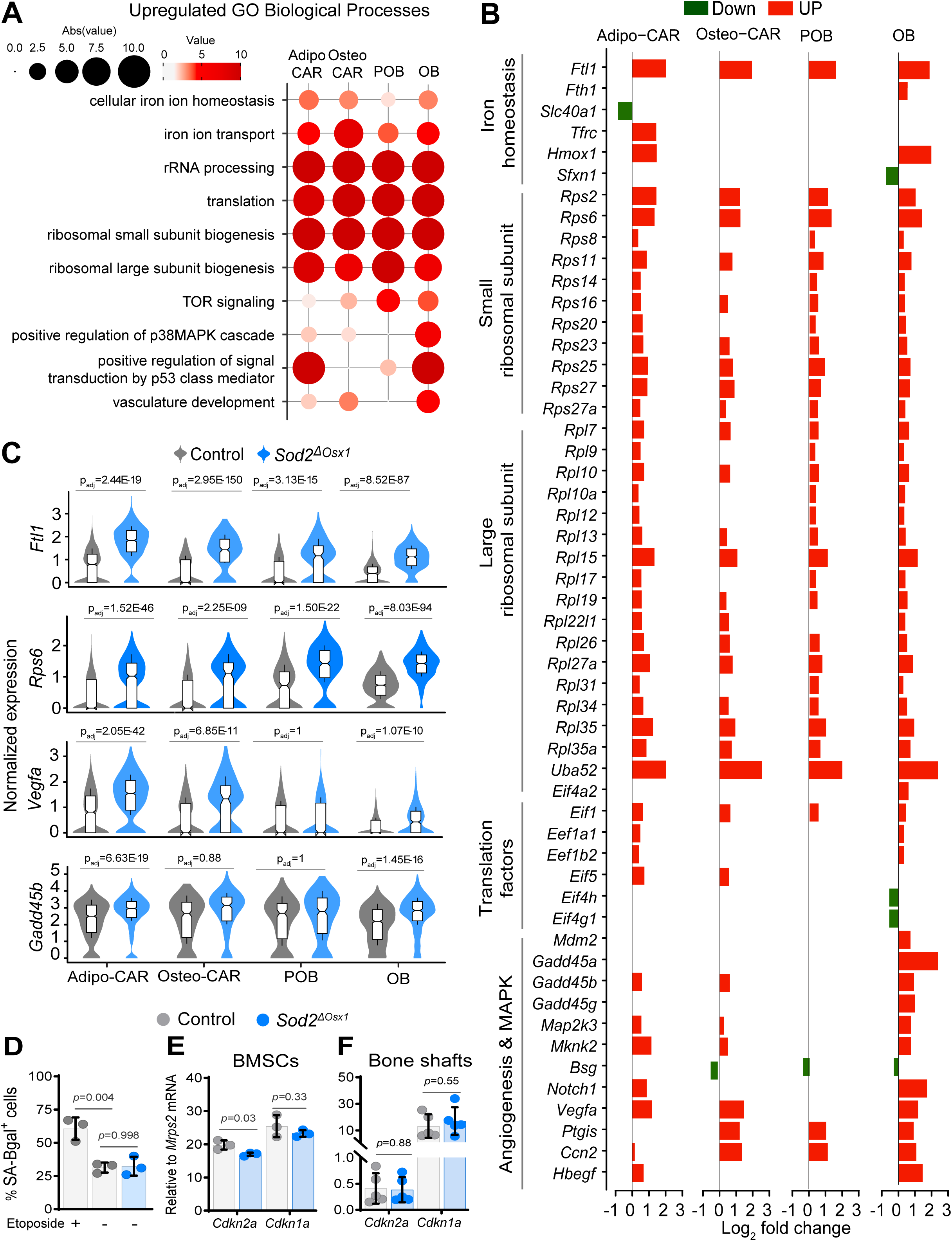
*Sod2* deletion alters iron homeostasis and translation. (A) Dot plot displaying significantly upregulated Gene Ontology (GO) Biological Processes (P-adj < 0.05, gene count > 5). Larger circle sizes and darker colors indicate higher significance. (B) Log_2_FC of representative DEGs grouped by function (Iron homeostasis, Small ribosomal subunit, Large ribosomal subunit, Translation factors, and Angiogenesis & MAPK) in Adipo-CAR, Osteo-CAR, POB, and OB clusters from *Sod2*^Δ*Osx1*^ compared to control. (C) Violin plots showing normalized expression of *Flt1*, *Rps6*, *Vegfa*, and *Gadd45b* in Adipo-CAR, Osteo-CAR, POB, and OB clusters from *Sod2*^Δ*Osx1*^ compared to control. Box-and-whisker plots (center line: median; box limits: 25th/75th percentiles; whiskers ±1.5× interquartile range) (D) Quantification of SA-β-gal–positive BMSCs after 4 days in osteogenic culture; 10 µM etoposide used as a positive control (technical replicates, n = 3/group). (E and F) mRNA expression of *Cdkn2a* (p16) and *Cdkn1a* (p21) measured by qRT-PCR. (E) BMSCs cultured for 7 days in osteogenic medium (technical replicates n = 3/group) and (F) bone shafts from 22-week-old mice (biological replicates n = 5/group). See also Figure S4. (D-F) Data are mean ± SD and *P* values by a two-tailed unpaired Student’s t-test.

ROS can promote senescence in osteoblastic cells *in vitro* (30) and *Sod2* deletion in mice can also cause cellular senescence (6). Nonetheless, in our scRNA-seq data, we found little evidence of cellular senescence. Of 119 senescence-related genes, compiled from SenMayo (31), MSigDB (32), and CellAge data sets (33), many of these genes were downregulated in the Adipo-CAR cluster, and only a few, such as *Cdkn1a*, *Cdkn2a*, bone morphogenetic protein 2 (*Bmp2*) and vascular endothelial growth factor A (*Vegfa*) were upregulated in the osteoblast cluster (**Fig. S4A**). To further examine whether *Sod2* deletion could cause cellular senescence, we cultured BMSCs from *Sod2*^Δ*Osx1*^ and *Osx1-Cre* control mice and found that the number of SA-β-gal-positive cells was not affected by *Sod2* deletion (**Fig. 5D**). In contrast, an increase in the number of SA-β-Gal^+^ cells was easily seen when control cells were cultured in the presence of the DNA damaging drug etoposide, used here as a positive control. We also examined the expression of *Cdkn1a* and *Cdkn2a*, common markers of cellular senescence. *Cdkn2a* mRNA was decreased, while *Cdkn1a* was not different between cells from *Sod2*^Δ*Osx1*^ and *Osx1-Cre* control mice (**Fig. 5E**). Finally, we examined expression of *Cdkn1a* and *Cdkn2a* in cortical bone shafts of femur, which contain osteocytes, osteoblast, and other mesenchymal lineage cells. An increase in *Cdkn2a* and other senescence markers is easily seen in bone shafts from old when compared to young wild-type mice (34, 35). However, we found no differences in expression of the two cell cycle inhibitors in bone from *Sod2*^Δ*Osx1*^ (**Fig. 5F**).

### *Sod2* deletion decreased expression of extracellular matrix and osteoblast differentiation genes

Deletion of *Sod2* downregulated cellular processes related to osteogenesis and extracellular matrix across all cell types (**Fig. 6A**). The majority of bone matrix proteins are secreted by osteoblasts. Nonetheless, expression of type I collagen (*Col1a1*), the primary structural protein of bone matrix, was downregulated in all clusters. Several other collagen genes were downregulated in osteoblasts from *Sod2*^Δ*Osx1*^ mice including *Col3a1, Col11a1, Coll11a2, Col5a1,* and *Col5a2* (**Fig. 6B**). In addition, expression of protein disulfide isomerases (PDIs), including *Pdia3, Pdia4, and Pdia6*, essential for disulfide bond formation during protein folding in the ER, was decreased. Both calnexin (*Canx*) and calreticulin (*Calr*), which are chaperones critical for the biosynthesis of glycoproteins in the ER, were down-regulated, as was the collagen-specific chaperone heat shock protein 47 (Hsp47), encoded by *Serpinh1*, which stabilizes triple-helical procollagen and prevents lateral aggregation. The small leucine-rich repeat proteoglycan (SLRP) family is a group of proteins and proteoglycans that are found in the extracellular matrix and regulate matrix assembly and cell signaling. In Pre-osteoblasts, Osteo-CAR and Adipo-CAR, expression of SLRP members such as osteonectin (*Sparc*), decorin (*Dcn*), lumican (*Lum*), and osteomodulin (*Omd*) were downregulated. Osteo-CAR cells are thought to contain a long-lived osteoblast precursor population (24, 36). Osteo-CAR cells from *Sod2*^Δ*Osx1*^ mice had decreased expression of osteoblast differentiation markers such as *Alpl, Bglap,* and distal-less homeobox 5 (*Dlx5*). These cell types also had decreased expression of the parathyroid hormone receptor (*Pth1r*) (**Fig. 6B, 6C**). Parathyroid hormone (PTH) is a major stimulator of bone formation (37). Together with the decreased mineralization shown in **Fig. 1F**, these data suggest that *Sod2* deletion decreases extracellular matrix (ECM) production and osteoblast differentiation.

**Figure 6.**
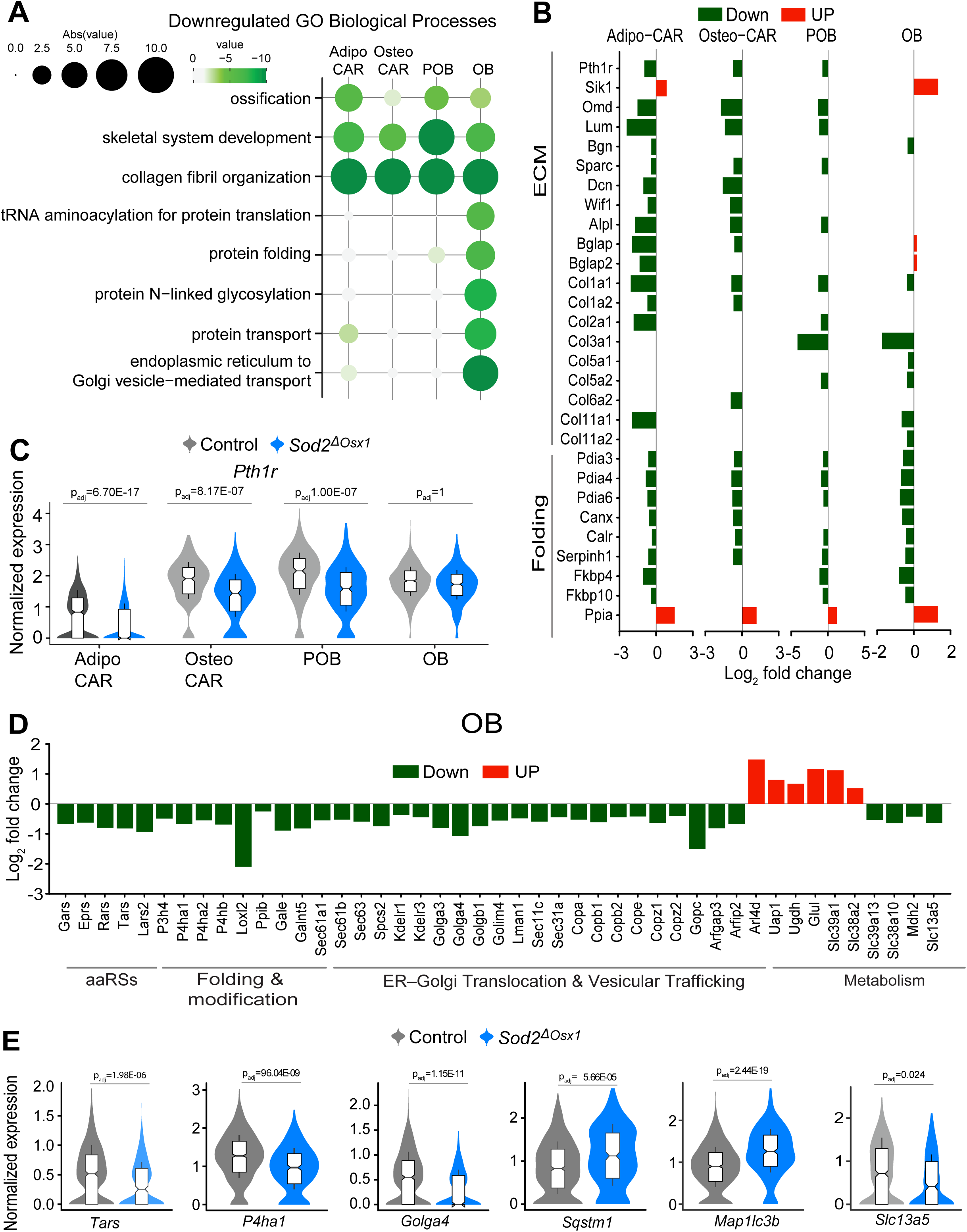
*Sod2* deletion alters markers of proteostasis in osteoblasts. (A) Dot plot displaying significantly downregulated Gene Ontology (GO) Biological Processes (P-adj < 0.05, gene count > 5) in Adipo-CAR, Osteo-CAR, POB, and OB clusters from *Sod2*^Δ*Osx1*^ compared to control. Larger circle sizes and darker colors indicate higher significance. (B) Log_2_FC of representative DEGs grouped by folding and extracellular matrix (ECM) in Adipo-CAR, Osteo-CAR, POB, and OB clusters. (C) Violin plots displaying the normalized expression of *Pth1r* in Adipo-CAR, Osteo-CAR, POB, and OB clusters from control and *Sod2*^Δ*Osx1*^ mice. (D) Log_2_FC of representative DEGs in OB clusters, grouped by aminoacyl-tRNA synthetases (aaRSs), protein folding and modification, ER-Golgi translocation and vesicular trafficking, and metabolism. (E) Violin plots of normalized expression of *Tars, P4ha1, Golga4, Sqstm1,* and *Map1lc3b* significantly altered by Sod2 deletion in OB. Data in (C and E) are box-and-whisker plots (center line: median; box limits: 25th/75th percentiles; whiskers ±1.5× interquartile range)

### Osteoblasts lacking *Sod2* have decreased expression of genes implicated in collagen processing

We also noted that osteoblasts had decreased gene markers for protein folding and transport including tRNA aminoacylation and protein N-linked glycosylation (**Fig. 6D, 6E**). Specifically, aminoacyl-tRNA synthetases (aaRS), such as *Eprs, Tars, Rars, Gars,* and *Lars2*, responsible for covalently linking codons to their corresponding amino acids were decreased. Several prolyl and lysyl hydroxylases such as *P3h4, P4ha1, P4ha2,* and *P4hb,* as well as lysyl oxidase *Loxl2,* which is critical for stabilizing the collagen triple helix, were decreased in osteoblasts lacking Sod2 (**Fig. 6D, 6E**). Components of the ER-to-Golgi transport machinery such as COPI complex subunits *Copa, Copb1, Copb2, Cope, Copg, Copz1,* and *Copz2,* as well as ER translocation components *Sec61a1, Sec61b, Sec63, Sec31c, Sec11c* were also downregulated. In line with a disruption in protein folding, the autophagy cargo markers p62 (*Sqstm1*) and microtubule-associated protein 1 light chain 3b (*Map1lc3b*) were upregulated, perhaps in an attempt to clear misfolded proteins (**Fig. 6E**). Several genes implicated in cellular metabolism were altered with Sod2 deletion. For example, the citrate transporter *Slc13a5*, which is highly expressed in osteoblasts(38), and the mitochondrial malate dehydrogenase (*Mdh2*) were downregulated, perhaps to avoid excessive accumulation of citrate due to inhibition of aconitase (**Fig. 6D, 6E**)

## Discussion

Cell type-specific knockout of *Sod2* in mice is a well-established approach for investigating specific effects of excessive mitochondrial ROS (39–41). We show that deletion of *Sod2* in *Osx1-Cre* targeted cells caused low bone mass, associated with a decreased osteogenic potential of osteoblast lineage cells, consistent with findings from previous studies using *Runx2-Cre* and *Dmp1-Cre* (6, 7). Importantly, our findings indicate that deletion of *Sod2* increases superoxide levels and reduces mitochondria respiration in osteoblastic cells mimicking some of the effects of aging. Deletion of *Sod2* also decreased the cellular levels of NAD^+^ and supplementation with NR improved OCR in osteoblastic cells cultured from *Sod2*^Δ*Osx1*^ mice. However, NR administration did not alter the low bone mass of *Sod2* deficient mice, indicating that a decrease in NAD^+^ is not a major mediator of the effects of mitochondrial ROS in bone cells. In contrast, administration of NR attenuates the loss of bone mass with aging(13). Most likely, mechanisms other than ROS are predominantly responsible for the decline in NAD^+^ with age.

Our scRNA-seq analysis of mesenchymal lineage cells from bone of *Sod2*^Δ*Osx1*^ mice shed light on target cells and potential mechanisms responsible for the deleterious effects of excessive ROS in osteoblastic cells *in vivo*. Osteoblast precursors such as osteo-CAR and pre-osteoblasts had decreased expression of osteoblastic genes, suggesting decreased differentiation potential. These findings are in line with previous work suggesting that ROS inhibits osteoblast differentiation via FoxO transcription factors (42) and by oxidizing and degrading RUNX2 (43). Interestingly, our single cell transcriptomic analysis revealed that osteoblast precursor cells from *Sod2*^Δ*Osx1*^ mice have lower expression of *Pth1r*. PTH is a critical regulator of blood calcium levels by promoting bone remodeling (37). PTH signals through PTH1r, present in osteoblast lineage cells to stimulate both bone resorption and bone formation. Adult mice lacking *PTH1r* in osteoblast lineage cells exhibit accelerated bone loss with age, suggesting that PTH signaling protects mice from age-related bone loss (44). Inhibition of salt-inducible kinases (SIKs) in osteoblasts and osteocytes is an important mediator of PTH actions on the skeleton (45, 46). In addition to the decrease in *Pth1r* expression, *Sod2*^Δ*Osx1*^ mice exhibited an increase in *Sik1* expression in osteoblasts. Together, these findings suggest that a decrease in osteoblastic cell response to endogenous PTH might contribute to the inhibition of osteogenesis by ROS.

Mitochondrial ROS can diffuse to the endoplasmic reticulum, alter its redox environment and disrupt protein processing (47–49). We identified changes in biological processes related to disrupted proteostasis in multiple cell clusters. Interestingly, osteoblasts were particularly impacted, most likely because of their major function of producing and secreting many bone matrix proteins. Collagen 1a, the primary structural protein of bone matrix, relies on a series of redox-dependent oxidative post-translational modifications for proper fibril formation and extracellular matrix integrity (50). For example, the formation of disulfide bridges during protein folding is catalyzed by PDI and entails a redox reaction that leads to H_2_O_2_ generation. In *Sod2* knockout osteoblasts, several genes involved in these post-translational modifications such as PDIs, hydroxylases, PPIases, and chaperones, were downregulated, perhaps to curb a further increase in ROS. These findings are in line with recent evidence that ROS in growth plate chondrocytes inhibits PDI-mediated disulfide bridge formation, oxidative protein folding, and ER proteostasis and negatively affects matrix collagen properties (51). Mice with connective tissue-specific *Sod2-*deficiecy also exhibited reduced collagen deposition and fibril thickness in the skin’s dermis (41). Accumulation of oxidative protein damage and a decline in proteostasis are closely linked to aging and age-related diseases (52, 53). Indeed, aged mice expressing a mitochondria-targeted catalase transgene (mCAT) maintain a youthful proteome in heart (54). We have shown that targeted overexpression of the mCAT transgene in the mesenchymal lineage attenuates age-related bone loss (5). The capacity of mCAT to sustain proteostasis likely mitigates age-related pathologies, including bone loss.

Mitochondrial ROS has been causally implicated in cellular senescence (55–57). Accordingly, *Sod2* deficiency induces DNA damage and promotes cellular senescence in some tissues (41, 57). Schoppa et al. have shown that deletion of *Sod2* in Runx2-expressing cells causes senescence which might contribute to the decreased bone formation (6). Nonetheless, we found no markers of senescence in cultured osteoblastic cells, freshly isolated mesenchymal cells, or cortical bone from *Sod2*^Δ*Osx1*^ mice. While the reasons for this inconsistency remain unknown, it is possible that the age of the mice might contribute to the different results. Indeed, Schoppa et al. (6) reported an increase in cellular senescence in bone of 12-month-old mice, while our analysis was conducted in younger mice. Thus, deletion of *Sod2* might accelerate bone cell aging and increase the number of senescent cells at a time when these remain low in control mice. Alternatively, *Runx2-Cre* might target mesenchymal cells that are not targeted by *Osx1-Cre*, and these might become senescent with *Sod2* deletion. In any case, because the decrease in bone mass with *Sod2* deletion is seen as early as 3-month-of-age, it is unlikely that cellular senescence represents a major contributor to the decrease in bone mass. Thus, mitochondrial ROS might not represent major inducers of osteoblastic cell senescence with aging. This assumption aligns with observations that attenuation of mitochondrial ROS does not prevent cellular senescence in adipose tissue of aged mice (58).

This study has limitations. In our mouse model of *Sod2* deletion, Osx1-Cre was active throughout development and growth in hypertrophic chondrocytes, osteoblasts, and other mesenchymal cells. Hence, subtle patterning effects, as well as the mixed background of the mice might have influenced the development of the phenotype. However, the low bone mass of the *Sod2*^Δ*Osx1*^ mice was similar to the one seen when *Sod2* was deleted with Dmp1-Cre or Runx2-Cre (6, 7). Dmp1-Cre does not target chondrocytes, and these previously described models have genetic backgrounds that are different from the *Sod2*^Δ*Osx1*^ mice. Together, these observations provide confidence that the low bone mass seen in our model was due to *Sod2* deletion in osteoblastic cells. We could not determine the cellular changes responsible for the low bone mass phenotype in 26-week-old mice. However, the scRNA-seq analysis performed with cells collected from 19-week-old *Sod2*^Δ*Osx1*^ mice suggests that the number of osteoblasts is decreased in line with findings from similar models (6, 7). Lastly, our scRNA-seq data reflect the long-term effects of mitochondrial ROS in cells of the mesenchymal lineage. Thus, it remains unclear how the physiological properties of osteoblastic cells are acutely altered by ROS. Future studies using Osx1-Cre or other models in which Cre activity can be temporally controlled should address this gap in knowledge.

Many aging-related pathologies, including loss of bone mass, are causally linked to ROS dysregulation. Nonetheless, the molecular mechanisms underlying these links are lacking. Work herein sheds light on oxidative stress-driven molecular changes in osteoblastic lineage cells. Our findings suggest that mitochondrial ROS dysregulates protein homeostasis, particularly in osteoblasts. Future research should elucidate how changes in proteostasis alter bone mass. Our work also indicates that the loss of bone caused by excessive ROS is not mediated by changes in NAD^+^ or cellular senescence, clarifying potential interconnections between these common mechanisms of aging. We propose that treatment combinations aimed at decreasing ROS, increasing NAD^+^, and eliminating senescent cells might confer additive effects in delaying osteoporosis and reducing fracture risk with aging.

## Experimental procedures

### Animal experiments

All animal procedures (protocol #AUP 4080(1)) were approved by the Institutional Animal Care and Use Committee (IACUC) of the University of Arkansas for Medical Sciences and conducted in compliance with the Public Health Service Policy, the NIH Guide for the Care and Use of Laboratory Animals (2011), the AVMA Animal Euthanasia Guidelines (2020), and the ARRIVE guidelines 2.0 (59). All experiments and methods were performed in accordance with these guidelines and regulations.

*Sod2* floxed (^f/f^) mice were obtained from cryopreserved sperm (provided by Lee Ann MacMillan-Crow at the University of Arkansas for Medical Sciences). Offspring from the *in vitro* fertilization (129/Sv and C57BL/6) were backcrossed to C57BL/6J mice (CD45.2) for four generations. Mice with conditional deletion of *Sod2* in osteoblast lineage cells were generated using a two-step breeding strategy. Hemizygous *Osx1-cre* transgenic mice (*B6.Cg-Tg(Sp7-tTA,tetO-EGFP/cre)^1Amc/J^;* Jackson Laboratory, Strain# 006361) were crossed with *Sod2^f/f^* mice to produce offspring heterozygous for the floxed Sod2 allele with (*Sod2^f/+^; Osx1-cre*) or without (*Sod2^f/+^)* the Cre allele. These mice were intercrossed to generate *Sod2*^Δ*Osx1*^ and littermate controls (*Osx1-cre*) mice. Young (6 months) and old female (24 months) C57BL/6J mice were obtained from the NIA-supported colony at Charles River Laboratories or purchased from Jackson Laboratory (Jackson Laboratory, Strain# 000664).

All mice were housed in groups of 2–5 animals per cage under controlled conditions (23°C, 60–70% humidity, 12-hour light/dark cycle), with *ad libitum* access to food and water. Breeder mice were fed the 5V5M diet (LabDiet, Cat. #3002906-704). All other mice were maintained on the 5V5R diet (LabDiet, Cat. #3002909-203). Body weight was measured before euthanasia. Mice were euthanized by CO₂ asphyxiation at designated time points, followed by cervical dislocation.

### Genotyping of transgenic mouse lines

All transgenic mouse colonies were genotyped by PCR using the REDExtract-N-Amp™ Tissue PCR Kit (Sigma-Aldrich, Cat# XNAT-100RXN) according to the manufacturer’s instructions. Briefly, offspring were tail-clipped at weaning (21 days) and before sacrifice. Genomic DNA was extracted by incubating tail samples in extraction buffer for 10 minutes at room temperature, followed by heating at 95°C for 3 minutes and neutralization. Please note that during genotyping of Sod2 floxed mice, we determined that the neomycin phosphotransferase (neo) selection cassette that was contained in the original conditional allele was missing, likely due to an unintended recombination event. Sequencing of PCR products produced using primers P1 (5′-CGA GGG GCA TCT AGT GGA GAA G-3′) and P2 (5′-TTA GGG CTC AGG TTT GTC CAT AA-3′), under the following PCR conditions 94°C −5min, 33x(94°C −35s, 58°C −35s, 72°C −35s), 72°C −10min, 12°C -end. The wild-type band is 500 bp, and the mutant band is 540 bp, revealing that the 540 bp product contained the loxP site predicted to exist after recombination of the loxP sites flanking the neo cassette. PCR amplification of *Osx1-cre* was performed using primers CREgeno fwd (GCGGTCTGGCAGTAAAAACTATC) and CREgeno rev: (GTG AAACAGCATTGCTGTCACTT), under the following PCR conditions 94°C - 3min, 35x(94°C −45s, 55°C −45s, 72°C −1min), 72°C −10min, °C -end. PCR products were separated on a 3% agarose gel and visualized using a ChemiDoc XRS+ Gel Imaging System (Bio-RAD).

### Nicotinamide riboside (NR) administration

Female *Sod2*^Δ*Osx1*^ and littermate control mice were randomized based on BMD into vehicle or nicotinamide riboside (NR) treatment groups. 12 mM NR (ChromaDex, Mfr Part # PHTN-SP001-30S-1) was administered in filtered drinking water, provided *ad libitum* in light-protected bottles, and replaced every day (except Sundays) for four months.

### Bone mineral density (BMD)

BMD was determined in mice using dual-energy X-ray absorptiometry (DEXA) with a PIXImus densitometer (GE Healthcare Lunar, software v2.0), as previously described (60). Each scan had an acquisition time of 4 minutes and an analysis time of 6 minutes. Mice were sedated with 2% isoflurane to ensure they remained motionless during the scans. Spinal BMD was assessed using the L1–L6 vertebrae, and femoral BMD was determined from the entire right femur. Quality control, performed with a proprietary skeletal phantom, yielded a mean coefficient of variation below 2%. Vital signs (righting reflex, respiration, and heart rate) were monitored during sedation to facilitate rapid post-examination recovery.

### Micro-computed tomography (μCT) scanning

Micro-computed tomography (μCT) was used to measure cortical and trabecular architecture of the fifth lumbar vertebra (L5) and right femur, as previously described (61). L5 vertebra and femurs were dissected, cleaned of soft tissues, fixed in 10% Millonig’s formalin (Leica Biosystems, Cat# 3800598) with 5% sucrose overnight, and gradually dehydrated in a graded series of ethanol (70%, 80%, 90%, and 100%) at 4°C. Dehydrated bones were then loaded into a 12.3-mm-diameter scanning tube and scanned by a µCT (Scanco Biomedical, model# µCT40) to generate three-dimensional voxel images (1024D×D1024 pixels) of the bone samples. A Gaussian filter (sigmaD=D0.8, supportD=D1) was used to reduce signal noise, and a threshold of 200 was applied to all scans at medium resolution (ED=D55DkVp, ID=D145DµA, integration timeD=D200Dms). Bone parameters were measured using Scanco Eval Program v.6.0. For vertebral analysis, the entire L5 vertebral body was scanned, and cortical bone and primary spongiosa were manually excluded from the measurements. Trabecular measurements were made by drawing contours every 10–20 slices and using voxel counting for bone volume per tissue volume and sphere-filling distance transformation indices without presumptions about the bone shape as a rod or plate for trabecular microarchitecture. For femoral analysis, scanning was performed from a point immediately distal to the third trochanter down to the proximal edge of the distal growth plate. Cortical dimensions at the diaphysis were determined by analyzing 18 slices centered at the midpoint of the bone length as determined in scout view. Calibration and quality control were performed weekly using five density standards, and spatial resolution was verified monthly using a tungsten wire rod. Beam hardening correction was based on the calibration records. Image acquisition and analysis protocols adhered to the guidelines of the Journal of Bone and Mineral Research (62).

### Bone histomorphometry analysis

Mice received intraperitoneal injections of calcein (10 mg/mL in 0.9% NaCl and 2% NaHCO₃; Millipore-Sigma) 7 and 3 days before harvest. Lumbar vertebrae (L2–L4) were dissected, cleared of soft tissue, and fixed overnight at 4 °C in 10 % Millonig’s formalin with 5 % sucrose (Leica Biosystems, Cat# 3800598). After graded ethanol dehydration (70 %, 80 %, 90 %, 100 %) at 4 °C, specimens were embedded in methyl methacrylate and sectioned at 5 µm thickness. Primary measurements included bone surface (BS), single-labeled surface (sLS), double-labeled surface (dLS), and interlabel thickness (Ir.L.Th). Derived indices—mineralizing surface (MS, %), mineral apposition rate (MAR, µm/day), and bone formation rate (BFR/BS, µm³/µm²/day)—were calculated according to ASBMR guidelines (63). All histology measurements were made in a blinded fashion. Fluorescent images were acquired using an Olympus BX53 microscope and analyzed with OsteoMeasure software (OsteoMetrics).

### Serum biochemistry

Blood was collected via retro-orbital bleeding and allowed to clot for 1 hour at room temperature. Samples were then centrifuged at 1,000 × g for 10 minutes, and serum was collected, aliquoted, and stored at –80°C. Serum levels of the bone remodeling markers P1NP (Immunodiagnostic Systems, Cat# AC-33F1) and CTX-1 (Immunodiagnostic Systems, Cat# AC-06F1) were quantified by ELISA according to the manufacturer’s instructions.

### Bone marrow-derived stromal cells (BMSCs)

Total bone marrow cells were obtained by flushing the femurs and tibiae with isolation medium composed of α-Minimum Essential Medium (α-MEM) supplemented with 20% fetal bovine serum (FBS; Gibco™, Cat# 10091155) and 1% penicillin/streptomycin (PS; Gibco™, Cat# 15140122) (30). The suspension was passed through a 70 μm cell strainer and centrifuged at 300 × g for 5 minutes. Red blood cells were lysed with ACK buffer (0.01 M EDTA, 0.011 M KHCO₃, 0.155 M NH₄Cl, pH 7.3) for 1 minute, followed by centrifugation at 300 × g for 5 minutes. Technical replicates were generated by pooling BMSCs from 3–6 mice/group. Cells were filtered again through a 70 μm strainer (Corning, Cat# 352350) and cultured in 10-cm dishes with α-MEM containing 20% FBS, 1% PS, and 50 μg/mL ascorbic acid (Sigma-Aldrich, Cat# A4403). Cultures were maintained at 37°C in a humidified atmosphere containing 5% CO₂ and the medium was replaced every 3 days. Adherent stromal cells were detached using 0.5% trypsin-EDTA (Gibco™, Cat# 15400054) and re-plated for osteogenic differentiation or other downstream (mtROS, mt ΔΨm, Seahorse, NAD^+^, SA-β-gal) assays.

### Osteogenic differentiation and alizarin red S staining

BMSCs were seeded in 12-well plates at a density of 2D×D10DDcells/well in a medium containing 10% FBS, 1% penicillin-streptomycin (PS), and 50 µg/mL ascorbic acid. After 2 days, osteogenic differentiation was induced by adding 10DmM β-glycerophosphate (Sigma-Aldrich, Cat# G9422), with media changes every 3 days.

On day 7 of osteogenic differentiation, total RNA was collected for RT-qPCR analysis of osteoblast-specific genes. Mineralization was assessed on day 21 using Alizarin Red S (AR-S) staining (Sigma-Aldrich, Cat# A5533). Briefly, cells were fixed in 10% Neutral buffered formalin for 30 minutes at room temperature. After two washes with double-distilled water (ddH₂O), cells were stained with 40 mM AR-S solution (pH 4.2) and incubated in the dark at room temperature for more than 20 minutes, followed by four washing steps with ddH_2_O. After drying, the AR-S was destained with 10% cetylpyridinium chloride in 10 mM sodium phosphate (pH 7.0) (Sigma-Aldrich, Cat# C0732). The AR-S was quantified by absorbance measured at 562 nm against a known AR-S standard.

### Cell proliferation assay

To evaluate BMSCs proliferation, a BrdU incorporation assay was performed using a Cell Proliferation ELISA kit (Roche, Cat# 1647229) according to the manufacturer’s instructions. Briefly, BMSCs were seeded at 4,000 cells per well in 96-well plates containing α-MEM supplemented with 10% FBS. At 60-70 % confluence, cells were incubated with 10 µM BrdU for 48 hours. The cells were then dried and fixed, and their DNA was denatured with FixDenat solution for 30 minutes at room temperature. Next, a peroxidase-conjugated mouse anti-BrdU monoclonal antibody was added and incubated for 90 minutes at room temperature. After washing, a tetramethylbenzidine substrate solution was applied for 15 minutes at room temperature, and absorbance was measured at 450–620 nm using a microplate reader. Mean data were expressed as a ratio of the control (untreated) cell proliferation.

### Quantitative RT–PCR

Right femoral and tibial bone shafts were prepared by removing bone ends, flushing marrow, and cleaning soft tissue and periosteum, as previously described (64). The prepared shafts were then flash-frozen in liquid nitrogen and pulverized using a multi-well tissue pulverizer (BioSpec Products, Cat# 59012MS). Total RNA was extracted from pulverized bone and cultured cells using TRIzol reagent (Thermo Fisher Scientific, Cat# 15596026). For cells, a PBS wash preceded RNA extraction. Quantitation and determination of the 260/280 ratio of the extracted RNA were determined by NanoDrop™ 2000 spectrophotometer (Thermo Fisher Scientific). cDNA synthesis was performed from 1 µg of total RNA using the High-Capacity Reverse Transcription Kit (Applied Biosystems, Cat# 4368813) on a PTC-200 Peltier Thermal Cycler (MJ Research). Real-time qPCR was carried out using TaqMan™ Gene Expression Master Mix (Applied Biosystems, Cat# 4304437), gene-specific TaqMan™ probes, and primers *Sod2* (Mm00449726_m1); *Cdkn1a* (Mm00432448_m1); *Cdkn2a* (Mm00494449_m1)), on a Quant Studio 3 PCR System (Applied Biosystems; RRID:SCR_020238). Expression levels were normalized to the housekeeping gene *Mrps2* (Mm00475528_m1) and expressed as fold change using the 2^^−ΔΔCt^ method (65).

### Mitochondrial ROS

Mitochondrial superoxide (•O2−) levels were determined using the mitochondrial-specific probe MitoSOX™ Red (Thermo Fisher Scientific, Cat# M36008). Briefly, BMSCs were seeded at 2.0 × 10D cells/well in the 96-well black-walled plate (Corning™, Cat# 3603) and cultured for 2 days in α-MEM supplemented with 10% FBS and 50 μg/mL ascorbic acid. Cells were washed twice with 1× PBS and incubated with 2.5 μM MitoSOX™ Red for 20 minutes at 37°C. Following two additional washes with PBS, fluorescence (Ex/Em = 510/580 nm) was measured using a Cytation 5 Cell Imaging Multi-Mode Reader (BioTek Instruments, Inc.; RRID:SCR_019732) to assess MitoSOX oxidation. MitoSOX intensity was normalized to nuclei counts using Hoechst 33342 (Thermo Fisher Scientific, Cat# H3570) staining.

### Mitochondrial membrane potential

Mitochondrial membrane potential (mtΔΨm) in BMSCs from control and *Sod2*^Δ*Osx1*^ was measured using the JC-1 Mitochondrial Membrane Potential Probe (Invitrogen™, Cat# T3168) according to the manufacturer’s protocol. Fluorescence (510/580 nm) was measured using a Cytation 5 Cell Imaging Multi-Mode Reader (BioTek Instruments, Inc.; RRID:SCR_019732). The ratio of JC-1 red to green fluorescence (Ex/Em ≈ 530/590 nm for red, 485/535 nm for green) was reported. MtΔΨm in BMSCs from young and old mice was measured using the tetramethylrhodamine (TMRM) probe (Invitrogen™, Cat# T668) according to the manufacturer’s protocol. Briefly, BMSCs were seeded at 5.0 × 10D cells/well in 96-well black-walled plates (Corning™, Cat#3 603) and cultured for 5 days in α-MEM with 50 μg/mL ascorbic acid. Following culture, cells were washed with PBS and incubated for 30 min at 37°C in assay medium (120 mM NaCl, 3.5 mM KCl, 5 mM NaHCO₃, 1.2 mM Na₂SO₄, 0.4 mM KH₂PO₄, 20 mM HEPES, 1.3 mM CaCl₂, 1.2 mM MgCl₂, 10 mM sodium pyruvate, pH 7.4) containing 100 nM TMRM. Fluorescence intensity (548/574 nm) was measured using a Cytation 5 Cell Imaging Multi-Mode Reader (BioTek Instruments, Inc.; RRID:SCR_019732) and normalized to cell counts using Hoechst 33342 staining.

### Seahorse analysis

Oxygen consumption rates (OCR) were measured using Mito Stress Kit (Agilent Technologies Cat#103015-100) and XFe96 Seahorse Analyzer (Agilent Technologies; RRID:SCR_019545) following the manufacturer’s protocol. BMSCs from control and *Sod2*^Δ*Osx1*^ mice were seeded at 4.0 × 10D cells/well in 96-well Seahorse plates (Agilent Technologies, Cat# 103794-100) and cultured for 2 days in α-MEM supplemented with 10% FBS and 50 μg/mL ascorbic acid. For nicotinamide riboside (NR) treatment, 5mM NR was added with a media change and cells were treated for an additional 16 hours. BMSCs from young and old mice were seeded at 5.0 × 10^4^ cells/well and cultured for 5 days in the same medium. Before the assay, cells were washed twice and incubated for 1 hour at 37°C (no CO₂) in 180 μl of Seahorse Base Medium (Agilent Technologies Cat# 103575) supplemented with 2 mM L-glutamine, 5.5 mM D-glucose, and 1 mM pyruvate. OCR was measured under basal conditions and after the sequential addition of 1.5 μM oligomycin, 2 μM FCCP, and 0.5 μM each of rotenone and antimycin A. Basal respiration was defined by the initial OCR values. Maximal respiration was calculated as FCCP-stimulated OCR minus OCR after rotenone/antimycin A treatment. Cells were stained with Hoechst 33342 (Thermo Fisher Scientific, Cat# H3570), counted using a Cytation 5 Cell Imaging Multi-Mode Reader (BioTek Instruments, Inc.; RRID:SCR_019732), and raw OCR values were normalized to the cell number. The data were analyzed using Wave Software (Agilent Technologies).

### NAD^+^ measurement

NAD^+^ levels were measured in lysates from both scWAT and BMSCs using the EnzyChrom™ NAD^+^/NADH Assay Kit (Bioassay Systems, Cat# EFND-100) per manufacturer’s instructions. Snap-frozen samples from control and *Sod2*^Δ*osx1*-cre^ mice were used directly for NAD^+^ assay. Protein levels were quantified using the Bio-Rad colorimetric protein assay. BMSCs from control and *Sod2*^Δ*osx1*-cre^ mice were seeded at a density of 2 × 10^4^ cells/well in 96-well white-walled plates (Corning™, Cat# 3917) and cultured for two days in α-MEM supplemented with 10% FBS and 50 μg/mL ascorbic acid.

### Senescence-Associated β-Galactosidase (SA-β-gal) assay

BMSCs were seeded in six-well plates and cultured for two days. The medium was then replaced, and cells were treated with B-glycerophosphate (Gibco™, Cat# G9422), while positive control groups received 10 μM etoposide (Calbiochem, Cat# 341205) for 3 days. Cells were washed with PBS and fixed with 4% paraformaldehyde (Sigma Aldrich, Cat# S0876) in PBS for 8 minutes, followed by two additional washes with PBS. For staining, the plates were incubated overnight (16–18 hours) at 37°C with a staining solution containing 1 mg/mL X-Gal (Sigma Aldrich, Cat# B4252), 40 mM citric acid/sodium phosphate buffer (pH 6.0), 5 mM potassium ferrocyanide (Calbiochem, Cat# 341205), 5 mM potassium ferricyanide (Sigma Aldrich, Cat# P3289), 150 mM sodium chloride, and 2 mM magnesium chloride (Fisher Scientific Cat# BP358). Following SA-β-gal staining, cells were counterstained with Hoechst dye (Thermo Fisher Scientific, Cat# H3570). Brightfield images were captured from at least three areas per well at 20× magnification using an inverted microscope (Olympus CKX41) equipped with a digital camera (Olympus DP20) and cellSens Standard software (Olympus Corp.). The same fields were imaged under the blue fluorescence channel to visualize Hoechst-stained nuclei. The proportion of SA-β-gal-positive cells was quantified using ImageJ software (NIH, RRID: SCR_002798)

### Isolation of endosteal mesenchymal cells for scRNA-seq

Mesenchymal cells associated with the endosteal and trabecular bone surfaces were isolated from the femurs and tibias of 4-month-old *Sod2*^Δ*Osx1*^ and littermate control mice (n=1 per group), as described previously (24, 35). After euthanasia, femurs and tibias were cleaned of soft tissues, and the periosteum was removed by scraping with a scalpel. The epiphyses were excised, and bone shafts were cut longitudinally. Bone marrow was flushed using PBS supplemented with 1% bovine serum albumin (BSA). The remaining bone fragments were then cut into smaller pieces of approximately 1 mm using a clean scalpel on a glass plate. Bone fragments were subjected to four Liberase™ (2 Wunsch units/mL in HBSS) and three EDTA (5 mM, 0.1% BSA in PBS without calcium or magnesium) digestions in a 6-well plate, alternating between the two solutions (20 min each, 37°C, 200 RPM). After each digestion, the cell-containing supernatant was collected and immediately placed on ice. Cells were then pelleted by centrifugation (300 x g, 10 min). The supernatants were carefully aspirated, leaving approximately 50-100 µL. Pellets were resuspended in 500 µL of sorting buffer (PBS, 0.5% BSA, 2 mM EDTA). Following the final digestion, all fractions were pooled, centrifuged, and resuspended in 100 µL of sorting buffer.

A two-step immunomagnetic depletion process using MACS column separation (Miltenyi Biote, Cat# 130-042-303) was used to enrich the mesenchymal cell population. First, a lineage cell depletion kit (Miltenyi Biotec, Cat# 130-090-858) was used to deplete hematopoietic and endothelial cells, following the manufacturer’s instructions. Briefly, cells were incubated with a biotin-conjugated antibody cocktail, washed, and then incubated with anti-biotin microbeads. The cell suspension was then passed through an LS column, and the flow-through (lineage-negative fraction) was collected. To further deplete residual CD45+, CD117+, and CD31+ cells, the following microbeads were used: CD45 (Miltenyi Biotec, Cat# 130-052-301), CD117 (Miltenyi Biotec, Cat# 130-091-224), and CD31 (Miltenyi Biotec, Cat# 130-097-418). The cell suspension was incubated with each microbead set, washed, and then passed through a second LS column. The flow-through, containing the enriched endosteal mesenchymal cell population, was collected and filtered (70 µm, Corning™ Cat# 352350). Cells were then pelleted (300 x g, 10 min) and resuspended in sorting buffer. Cells were then pelleted (300 x g, 10 min) and resuspended in sorting buffer. Cells were immediately transported at 4°C to the UAMS Genomics Core for single-cell RNA sequencing.

### Single-cell RNA library preparation and sequencing

Single cells were stained with ReadyProbes Cell Viability Imaging Kit, Blue/Green (Thermo Fisher Scientific, Cat# R37609) and counted manually using a hemocytometer under an EVOS M7000 microscope (Thermo Fisher Scientific). Following cell counting, cells from each condition were encapsulated using a Chromium Controller (10X Genomics), and libraries were constructed using a Chromium Single Cell 3′ Reagent Kit (10X Genomics, PN-1000128) by the UAMS Genomics Core. Libraries were sequenced on an Illumina NovaSeq 6000 machine (RRID:SCR_016387) to generate FASTQ files.

### Single-cell RNA-seq data processing and analysis

The bioinformatics analysis of the scRNA-seq was followed our previous analysis of mesenchymal cells (24). The FASTQ files were then preprocessed using Cell Ranger software version 7.2 (10X Genomics, RRID:SCR_017344) to produce feature-barcode matrices, with alignments conducted using the mouse reference genome mm10. These feature-barcode matrices were imported for further analysis into R software (RRID:SCR_016341), utilizing the Seurat package version 5 (RRID:SCR_016341) (66–68). Cells containing between 1000 and 5,000 transcripts and mitochondria read content less than 15% were included in the analysis. The samples were integrated to minimize batch effect using canonical correlation analysis (CCA) analysis is method with to 50 principal components, The cell cluster analysis was performed based on shared nearest neighbor modularity optimization based clustering algorithm of Louvain algorithm with multilevel refinement. Uniform Manifold Approximation and Projection (UMAP) dimensional reduction technique was performed for visualization of the results. Gene-specific markers for individual clusters and differentially expressed genes (DEGs) were identified using the MAST algorithm (RRID:SCR_016340), which has shown favorable results in recent benchmarks (24). We used PIANO software (RRID:SCR_003200) to identify biological pathways and processes enriched in our sets of DEGs(69).

### Statistical analysis and visualization

The statistical tests performed, number of replicates, and error measures for each experiment are indicated in the figure legends. All *in vitro* assays were repeated three times. Statistical analysis relating to scRNA-seq was performed in R software. Other statistical analyses were conducted in GraphPad Prism 10, following verification of data adherence to normality, equal variance, and independent sampling assumptions. Significance was defined as *p*_adj_ ≤ 0.05 for GO terms and DEGs, and adjusted *p* ≤ 0.05 for all other tests. Sample sizes for animal experiments were based on prior studies, and no data were excluded from analysis. Figures were prepared using BioRender.com, Adobe Illustrator, and GraphPad Prism 10.

## Data availability

The scRNA-seq datasets generated and/or analyzed during the current study are available in the NCBI Gene Expression Omnibus (GEO) repository under BioProject number PRJNA1231044. This paper does not report the original code. All other data generated or analyzed during this study are included in this published article and its supplementary information files.

## Supporting information

This article contains supporting information.

## Conflict of interest

The authors declare that they have no known competing financial interests or personal relationships that could have appeared to influence the work reported in this paper.

## Acknowledgments

The authors would like to thank Lee Ann MacMillan-Crow, Ph.D., for generously providing the cryopreserved *Sod2 floxed (^f/f^)* mice and the CMDR Genetic Models Core for their assistance in retrieving them. We are grateful to the staff of the Bone Histology and Imaging Core and the UAMS Department of Laboratory Animal Medicine for their technical support. We also thank Olivia Reyes-Castro and Kimberly Richardson for their assistance with tissue collection.

## Author contributions

M.A. conceived the study. M.M.A., C.A.O., and M.A. designed experiments and analyzed the results. Q.F. recovered the *Sod2 floxed (^f/f^)* mice from cryopreserved sperm. M.M.A. conducted the breeding of conditional *Sod2* knockout mice. M.M.A., A.R.C., and A.W. performed nicotinamide riboside (NR) administration studies. M.M.A. performed micro-CT measurements. M.M.A. and A.M.C. performed *in vitro* studies. H.N.K. provided technical support and discussed the results. C.A.O. collected and processed samples for scRNA-seq. I.N., M.M.A., and M.A. analyzed the scRNA-seq data. M.M.A. and M.A. wrote the manuscript. All authors reviewed the manuscript.

## Funding and additional information

This work was supported by the US National Institutes of Health (R01AG068449, R01AR56679), Center for Musculoskeletal Disease Research COBRE (P20GM125503), and the UAMS Bone and Joint Initiative.

## Abbreviations

BMSCs: Bone marrow-derived stromal cells
CAR: Cxcl12 abundant reticular
DEGs: differentially expressed genes
GO: gene ontology
mtROS: mitochondrial reactive oxygen species
NAD⁺: nicotinamide adenine dinucleotide, oxidized
OCR: oxygen consumption rate
SASP: senescence-associated secretory phenotype
SA-β-gal: senescence-associated beta-galactosidase
scRNA-seq: single-cell RNA sequencing
SD: standard deviation
UMAP: uniform manifold approximation and projection
μCT: micro-computed tomography
ΔΨm: mitochondrial membrane potential

## Notes

### Competing Interest Statement

The authors have declared no competing interest.

### Summary of Updates

Figure 1 has been revised, exact replicate details have been updated in the figure legends, a paragraph on study limitations has been added to the Discussion section, and the supporting information has been updated accordingly

