## Supporting Figures S1-S5 for "Mechanisms of mitochondrial reactive oxygen species action in bone mesenchymal cells"

**Includes:**

**Figures S1-S5**

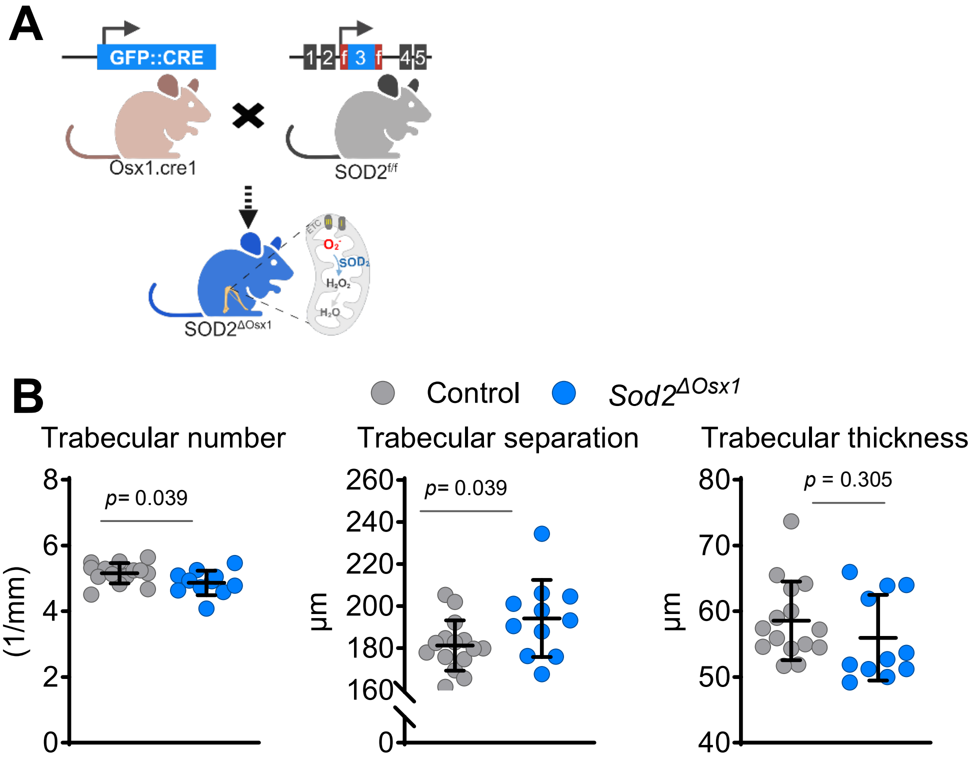


**Figure S1. Breeding strategy and μCT analysis of the fifth lumbar vertebrae from 26-week-old male *Sod2*$^{\Delta Osx1}$ mice**

(A) Breeding scheme for osteoblastic lineage-specific *Sod2* knockout mice (*Sod2*$^{\Delta Osx1}$), driven by the *Osx1* promoter. Hemizygous Osx1-cre transgenic mice were crossed with *Sod2*$^{f/f}$ mice. Resulting offspring, heterozygous for the floxed Sod2 allele with (*Sod2*$^{f/+}$*; Osx1-cre*) or without (*Sod2*$^{f/+}$) the Cre allele, were intercrossed to generate Sod2$^{\Delta Osx1}$ and littermate controls (*Osx1-cre*).

(B) Quantitative μCT analysis of trabecular parameters (number, separation, and thickness) in the fifth lumbar vertebra (L5) of control and *Sod2*$^{\Delta Osx1}$ mice (n = 9-10 mice/group).

Line and error bars represent mean ± S.D. P values by a two-tailed unpaired Student's t-test. **Related to Figure 1.**

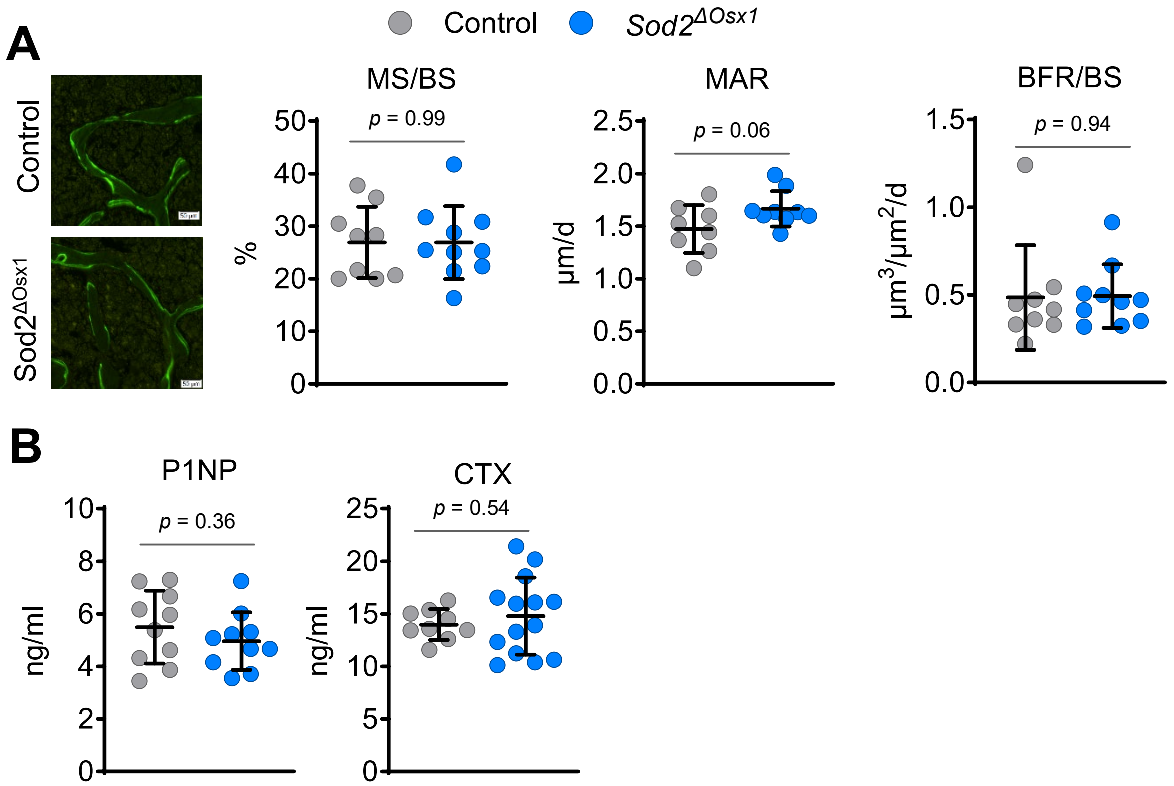


**Figure S2. Vertebral Histomorphometry analysis and serum bone turnover markers from 26-week-old male *Sod2*$^{\Delta Osx1}$ mice**

(A) Representative images of calcein double-labeling of trabecular bone from the 3rd lumbar vertebra, with quantification of mineralizing surface per bone surface (MS/BS), mineral apposition rate (MAR), and bone formation rate per bone surface (BFR/BS) (n = 9-10 mice/group). Scale bar: 50 µm.

(B) ELISA analysis of the serum N-terminal propeptide of type I procollagen (PINP) (n = 10 mice/group), and C-terminal telopeptide of type I collagen (CTX) (n = 9-14 mice/group).

Line and error bars represent mean ± S.D. P values by a two-tailed unpaired Student's t-test.. **Related to Figure 1**

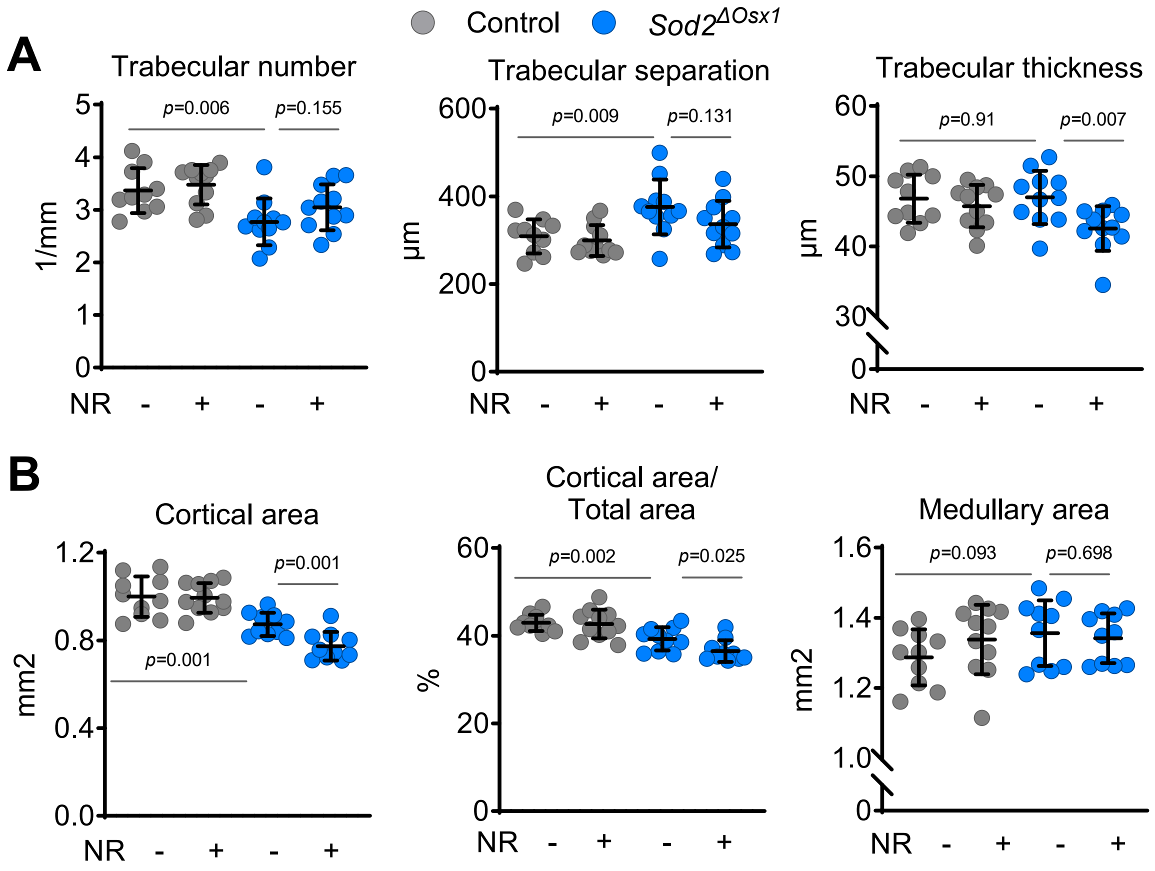


**Figure S3. μCT analysis of vertebral and femoral bone from 52-week-old female *Sod2*$^{\Delta Osx1}$ mice supplemented with NR**

(A) Quantitative μCT analysis of trabecular parameters (number, separation, and thickness) in the fifth lumbar vertebra (L5) (n = 9-10 mice/group)

(B) Quantitative μCT analysis of cortical area, cortical area/total area, and medullary area in the femur midshaft (n = 9-10 mice/group).

Line and error bars represent mean ± S.D. P-values by two-way ANOVA with Tukey's multiple comparisons test. **Related to Figure 3**

**A**

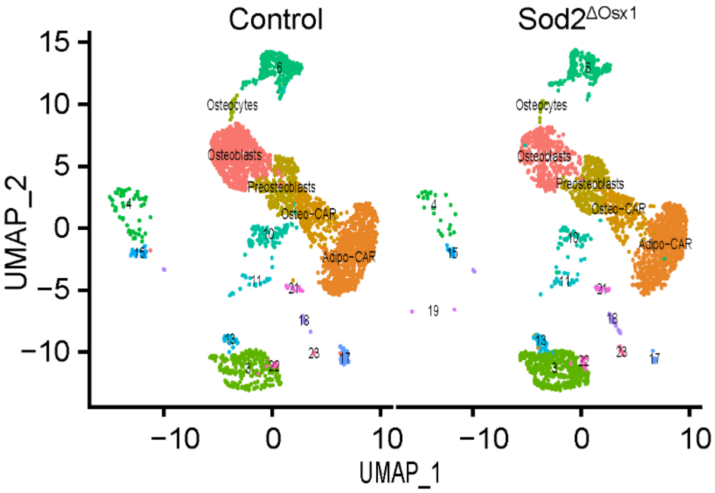


**Figure S4. UMAP projection of 15,224 endosteal cells**

(A) UMAP plot of cells isolated from the endosteal compartment, with non-mesenchymal cell clusters labeled numerically. **Related to Figure 4**

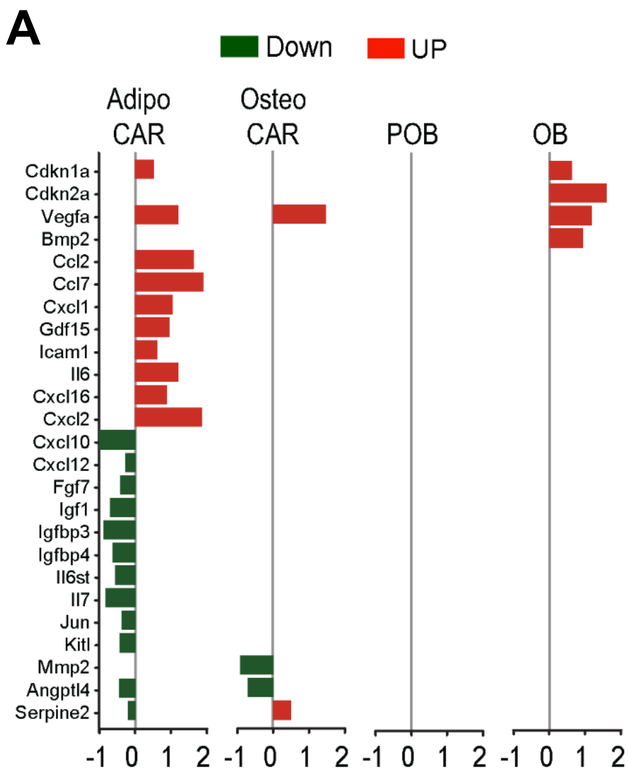


**Figure S5. Differentially expressed senescence-related genes in mesenchymal cells from *Sod2*$^{\Delta Osx1}$ and control mice**

(A) Log2FC values of 119 senescence-related genes, curated from SenMayo, MSigDB, and CellAge, that are differentially expressed between *Sod2*$^{\Delta Osx1}$ and control mice in the Adipo-CAR, Osteo-CAR, preosteoblast (POB), and osteoblast (OB) clusters. **Related to Figure 5**
